# Using an agent based model to inform sampling design for animal social network analysis

**DOI:** 10.1101/2024.05.25.595870

**Authors:** Prabhleen Kaur, Simone Ciuti, Michael Salter-Townshend, Damien Farine

## Abstract

Producing accurate and reliable inference from animal social network analysis depends on the sampling strategy during data collection. An increasing number of studies now use large-scale deployment of GPS tags to collect data on social behaviour. However, these can rarely capture whole populations or sample at very high frequencies. To date, little guidance exists when making *prior* decisions about how to maximise sampling effort to ensure that the data collected can be used to construct reliable social networks. We use a simulation-based approach to generate a functional understanding of how the accuracy of various network metrics is affected by decisions along three fundamental axes of sampling effort: coverage, frequency and duration. Researchers often face trade-offs between these three sampling axes, for example due to resource limitations, and thus we identify which axes need to be prioritised as well as the effectiveness of different deployment and analytical strategies. We found that the sampling level across the three axes has different consequences depending on the social network metrics that are estimated. For example, global metrics are more sensitive than local metrics to the proportion of the population tracked, and that among local metrics some are more sensitive to sampling duration than others. Our research demonstrates the importance of establishing an optimal sampling configuration for drawing relevant and robust inferences, and presents a range of practical advice for designing GPS based sampling strategies in accordance with the research objectives.

## Introduction

Social network analysis is a valuable tool for animal studies, providing insights into the structure, dynamics, and functioning of social relationships within the animal communities (***Croft et al., 2008***; ***Farine and Whitehead, 2015***; ***Krause et al., 2007***; ***Shizuka et al., 2021***; ***Sosa et al., 2021***). A challenge for studies using SNA is the ability to collect sufficient data to construct the network (***Kaur et al., 2023***; ***Smith et al., 2022***; ***Hoppitt and Farine, 2018***; Silk, 2018; ***Gilbertson et al., 2021***). Collat\ing social data ideally involves observing *many* or *all* individuals in a group or population at once. However, the cost and effort required to do so, especially in free-ranging populations, is often prohibitive (***Jung et al., 2018***; ***Hebblewhite and Haydon, 2010***; ***Krause et al., 2013***). Insufficient spatial coverage or biased sampling locations can lead to incomplete, inaccurate, or biased network representations (***Smith and Moody, 2013***; ***Davis et al., 2018***; ***James et al., 2009***). The duration and intensity of sampling efforts also impact the completeness and reliability of social network data, as short observation periods may underestimate overall network connectivity and dynamics (***He et al., 2022***; ***Fründ et al., 2015***; ***Lusseau et al., 2008***; ***Perreault, 2010***; ***Cross et al., 2012***). Moreover, biases in sampling, such as preferentially observing certain individuals or groups over others, provide an inaccurate picture of social interactions and network structure, affecting the reliability of inferences obtained from the data (***Espín-Noboa et al., 2018***; ***Davis et al., 2018***; ***Voelkl et al., 2011***; ***Cross et al., 2012***). While these potential problems are well established, previous work provides little guidance on how to overcome sampling limitations. Yet, there are likely to be some general principles that can be followed (***He et al., 2022***).

Most studies face trade-offs when determining the most appropriate sampling effort. These include sampling more broadly to increase the proportion of the population sampled versus sampling fewer individuals more often (***Davis et al., 2018***; ***Silk et al., 2015***; ***Smith and Moody, 2013***; ***Wey et al., 2008***). This issue is increasingly resolved by using automated tracking methods that can substantially increase the capture of data at the individual level (***Alarcón-Nieto et al., 2018***; ***Aplin et al., 2015***; ***Strandburg-Peshkin et al., 2015***). In wild populations, telemetry methods, in particlular GPS tracking, are becoming increasingly used to study social behaviours (***He et al., 2022***). However, GPS tracking also involves a number of trade-offs. For example, tags can increase sampling frequency (more points per unit of time) at the cost of sampling duration. Further, because of their high cost, decisions also need to be made about which individuals to deploy GPS trackers onto. Here again there are trade-offs, such as between sampling more intensively within a focal subset of the population versus spreading GPS devices more evenly across the population (Smith and Morgan, 2016; ***Smith and Moody, 2013***).

An issue for empiricists is that realistic sampling recommendations for wildlife contact networks are lacking, and little research has explored the impact of telemetry location frequency on network estimation (***Cross et al., 2012***; ***Gilbertson et al., 2021***; ***Costenbader and Valente, 2003***). One challenge with generating guidelines for how to sample wild populations is that we typically do not have a real baseline against which to evaluate our dataset. One approach has been to subsample large and intensively collected data (e.g. ***Davis et al. (2018)***). However, such datasets are not yet available for species where sampling challenges are most exacerbated, notably in species where there is continuous formation and dissolution of social groups (i.e. fisson-fusion dynamic, ***Aureli et al. (2008)***). One solution is to simulate population-level data, from which we can test different sampling strategies. ***Gilbertson et al. (2021)*** used simulations of individual movement trajectories to outline best practices for telemetry sampling, in particular for network studies of infectious disease in wildlife. ***Gilbertson et al. (2021)*** suggested prioritizing local network metrics and increasing sampling frequency to enhance accuracy, particularly in situations of low sampling coverage. However, it is unclear how general these findings are across, and whether they are influenced by nonrandom or geographically biased telemetry sampling. Additionally, the simulations assumed that individuals move independently. Yet we know that many animals also make social movements, which could also affect network inference. Further investigation is needed to understand sampling strategies and how association behaviours among individuals affect networks inferred using telemetry methods.

In this paper, we developed and used an agent-based model that incorporates both social and non-social factors affecting animal movements and interactions. Our objective was to understand the trade-offs associated with collecting empirical data using animal telemetry. Researchers frequently face trade-offs when allocating GPS resources for data collection, including decisions on the number of animals to monitor, the temporal resolution of observations, and the duration of monitoring required to gather reliable data aligned with research objectives. Here we focused on the accuracy of social network metrics estimates when moving along the three axes of sampling efforts, specifically (***He et al., 2022***; ***Gilbertson et al., 2021***):

- Sampling coverage : The number and allocation of GPS devices among individuals in one or more social groups.
- Sampling frequency : The temporal resolution at which GPS devices record data.
- Sampling duration : The total amount of time over which devices collect data.

We first quantified how increasing sample coverage, frequency, and duration contribute to a reduction of errors in different network metrics, spanning from the individual to the population. Next, we examined more directly the trade-off between sampling more intensively versus sampling for a longer duration. This allowed us to determine which sampling approach delivers the best estimations for a given amount of data. Finally, we examined the effect o different decisions with regards to sampling coverage. Specifically, we tested how selecting individuals that were more densely connected versus randomly selecting individuals across the broader population affected conclusions about the population network structure (***He et al., 2022***). We achieve these three tasks with the help of a custom agent based model designed in software R (R Core Team, 2022). Our models were inspired by ungulate movement, thus allowing us to portray the intricacies of the data collection process and answer such questions for this well-studied group. However, our results are also applicable to all species that live in societies with fission-fusion dynamics.

## Materials and Methods

### Model Scope and Conceptual Design

Our model structure follows the ODD criteria proposed by ***Grimm et al. (2020)*** for agent-based models. The model presented here is based on some fundamental assumptions that have allowed us to simplify the representation of complex animal decisions while capturing vital aspects of the movement process, and thus the drivers of encounters among individuals. See Section Model assumptions in the supplementary material for the assumptions of this agent based model.

### Environment Representation

We simulated an environment in the form of a grid structure. Figure 1 depicts the spatial organisation of the environment consisting of a two-dimensional 40 by 40 grid. Each cell in the grid represented a location where agents occur and denoted a distance unit (1 unit = 10 metres). The total area represented by this 40 by 40 grid is 16 hectares. The environment is also heterogeneous, with resource density varying across the grid. Each cell in the grid had a resource quality index value, which ranged from 0 to 1, with 0 indicating no availability of resources and 1 indicating maximum resource amount.

### Agent definition

Our model consisted of 500 individuals, or agents (Figure 1). An individual’s initial state could be one of two: foraging or resting. Each individual could be foraging or resting with a probability of 0.7 and 0.3. If an individual was in a foraging mode, they walked one unit each time point across the grid. If they were resting, remained in the same cell until their state changed. Each individual was also given an energy level at the start of the simulation. An individual’s energy level could range from 0 to 1, with 0 indicating the lowest energy level and 1 indicating the highest level of energy possible. We also defined connections among the individuals within the population, resulting in 16 sets of individuals that exhibited social preferences towards each other. The size of these social sets ranged from 19 to 41 animals. In each social set, a pair was assigned a connection with a probability of 0.9.

### Behavioural Rules

#### Resting state

If an individual was in resting state, they remained in the same cell and there was no movement. Their energy in the resting state lowered by 0.05 units per unit time.

**Figure 1.**
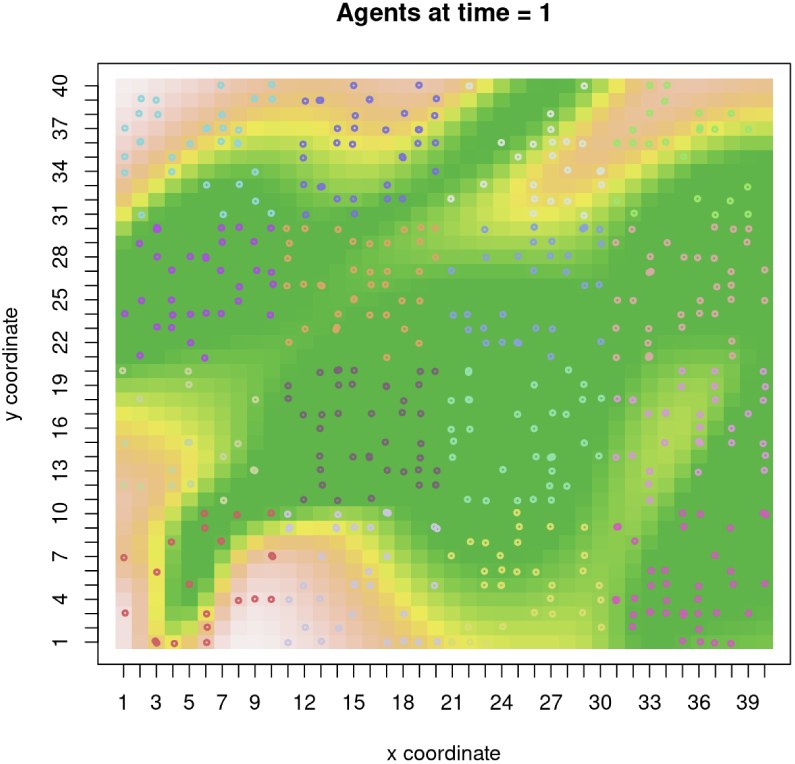
Simulated environment and agents. The simulated environment consists of a 40 x 40 grid cells. At time point 1, individuals’ locations are randomly distributed. The color of a grid cell represents it’s resource quality. The greener the cell, the better the resource available at that cell.

**Figure 2.**
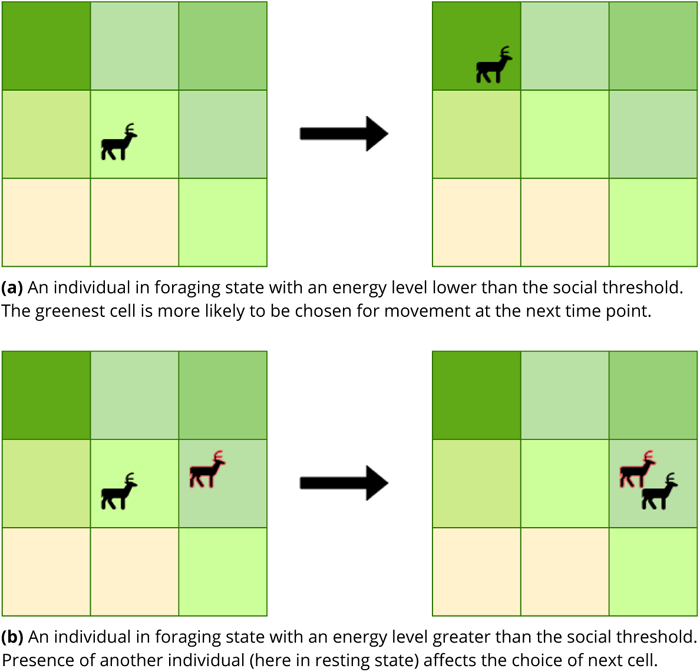
An individual’s movement decisions in absence and presence of its connection in a neighbouring cell.

#### Foraging state

At each time point, an individual in the foraging state moved to one of the nine immediate surrounding cells (including the current cell) as shown in Figure 2a. An individual’s movement and the choice of next cell was based on their current energy level and the presence of socially preferred conspecifics in the neighbouring cells.

An individual could opt to make social movements if one of their preferred conspecifics was present in a neighbouring cell or to go to a cell with a higher resource quality index. This decision was made based on their current level of energy and the social threshold. We defined the social threshold (set to 0.5) as the threshold below which individuals made movements exclusively based on food resources. Above this threshold, individuals moved preferentially towards strong social contacts. When making social movements, individuals’ energy decreased by 0.1 units per time point. When foraging, individuals either consumed resources or searched to find better quality food, depending on the habitat quality in the current cell and on their current energy level. If the current energy level was lower than the cell’s resource quality index, the individual used the resource and gained energy, otherwise they searched for a better resource (reducing their energy decreased by 0.1 unit per time point). Finally, foraging reduced the resource quality index of each cell, which was depleted following an exponential function of the current resource quality. The depletion function is given by:

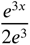

where *x* is the current resource quality index of the cell. The amount of resource quality depleted by an individual’s foraging was equal to the amount of energy acquired by that individual through resource consumption. See section Decision making process of an individual for a flowchart summary of the decision making process of an individual in the model.

In case, where multiple connections were present in different neighbouring cells, the decision was based on a two-step process. See Agent’s movement when multiple connections are present in multiple neighbouring cells for more details on this process.

### Parameter Settings

For an agent based model to be realistic, a good set of parameters is necessary to demonstrate the target behavior in a realistic way (***Tan and Bora, 2019***). In this model, there were several parameters that were needed to be tuned such that the model would reflect the nuances of the environment and social effects on movement. The model was calibrated over multiple set of parameters. For details, see section Model calibration in supplementary materials.

### Model Implementation And Data Collection

After setting up the grid, we performed the simulations for 4380 time points. Each time point in the simulation spanned two hours. This would depict the movement of animals recorded throughout a year, with observations made every two hours. The dataset consisted of 4380 time points, with each time point containing information on individual positions on the grid (given as (x,y) coordinates), their energy level, and their activity state (either foraging or resting).

As the goal is to understand how sampling levels affect the network metric accuracy, we sampled at different values spanning the three fundamental axes of sampling effort - sampling coverage, sampling frequency and sampling duration (Figure 3). The values of sampling coverage ranged from 25% to 100% where we analysed the networks obtained from a random selection of 125, 250, 375 and 500 individuals. For sampling duration, we analysed the data collected over 91 days, 182 days, 274 days and 365 days. This represented one quarter, one half, three quarters, and an entire year of observation. In terms of sampling frequency, networks were generated by recording the observations at different fixed intervals. The length of fixed intervals between subsequent observations were 2 hours, 4 hours, 8 hours and 16 hours. This lead to a sampling frequency of 12 fixes, 6 fixes, 3 fixes and 1.5 fixes per day. In this way, 64 samples were selected to obtain 64 network structures.

In a similar way, for the third aspect of the analysis, we sampled individuals with a coverage of 25% and 50 % but whole sets of individuals were sampled. For example, for a sampling coverage of 25%, 125 individuals were chosen belonging to just 5 groups as compared to all 16 groups in random sampling.

### Consideration Of GPS Error In Data Collection

In the process of gathering empirical data through GPS fixes, various sources of positional error may arise (***He et al., 2022***). To ensure greater realism in our simulated model, we incorporated these nuances into the data collection procedure. We used empirical data gathered by ***He et al. (2022)*** and generated a distribution of error magnitudes while taking a GPS fix. We then recorded location of each individual at each time point with a possibility of error. The error magnitude for each observation was sampled from the distribution of errors from obtained from the empirical data.

**Figure 3.**
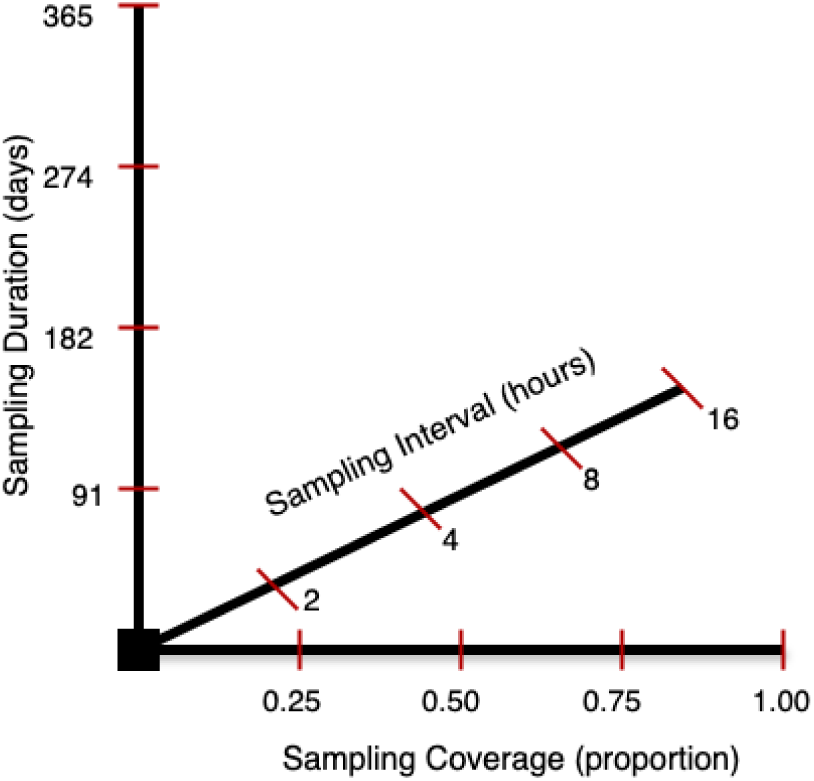
Sampling points along the three axes of sampling. Sampling along these points produced 64 networks. The sample obtained corresponding to the point (1.00, 2, 365) produced the network with highest accuracy.

In our simulation, as each cell in the grid comprised of a 10m X 10m region, an error value of magnitude less than 10m did not displace the individual. However, if an error value was between 10m and 20m, the recorded location would shift by one cell randomly in any direction and so on. Therefore, for each recording location of an individual in the simulation, there was a possibility of it’s position being wrongly recorded.

### Analysis

We obtained 64 network structures from the 64 samples that were generated as described above. We analysed these networks with respect to three global and three individual-level network metrics. The global network metrics used for analysis were edge density, assortativity, and transitivity and the individual-level metrics used for analysis were degree, strength, and betweenness.

Edge density (binary) is the ratio of the number of edges present to the maximum number of edges that the network can contain. Assortativity measures preference for the nodes of a network to connect to other nodes that are similar in some way (***Farine, 2014***). In this simulation, we had initialised individuals belonging to one of the 16 pre-defined sets of strongly connected individuals. Therefore, assortativity provided a measure of tendency of individuals were in the same location (i.e. inferred to be associated) with their preferential associates versus other individuals. We calculated assortativity using the assortnet package (***Farine, 2016***) in R. The assortativity value for the fully observed network was 0.8541. Transitivity of a network measures the probability that the connections of a node are also connected with each other.

The correlation analysis included three individual-level network metrics: degree, strength, and betweenness. The degree of a node represents the number of connections it has in the network. The strength of a node is equal to the sum of all the edge weights connecting to it. It is often referred to as weighted degree. The betweenness of a node is the number of times it appears on the shortest path between two other nodes in the network. Nodes with high betweenness values act as bridges connecting different parts of a network. Nodes in a network can be ranked according to their values of degree, strength, or betweenness. The agent base model was designed in R (R Core Team, 2022) where all analyses were carried out. We used the R package aniSNA (***Kaur, 2024***) to easily compute network metrics and analyse network structures.

### Accuracy

Our goal was to quantify three factors that impact accuracy. First, we looked at how the accuracy of projected network metrics changes as we moved along the three sample axes. We calculated the absolute relative error between the estimated values of the global network metrics and the observed value acquired from the full network as it provides a standardized measure that is independent of the scale or magnitude of the network metrics being compared. Full network refers to the network obtained from the fully simulated population and corresponds to the point (1.00, 2, 365) on the 3 dimensional sampling space (Figure 3). Absolute relative error is defined as the absolute difference between the true value and the estimated value, normalized by the true value and is calculated as follows:

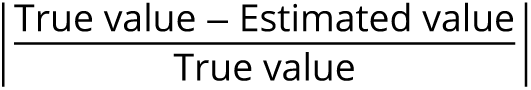

To determine the accuracy and rank preservation with respect to individual-level network metrics, we calculated the correlation coefficients between the network metric values of the sampled networks and the fully observed network. This analysis allowed us to acquire an understanding of how well the original ranks of the nodes were preserved in the sampled networks.

Secondly, we considered the trade-offs between sampling duration and frequency, as one often comes directly at the cost of the other. At each of the 16 levels of sampling (4 for sampling duration times 4 for sampling frequency), we calculated the total number of recorded observations as follows:

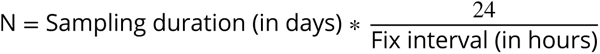

where,

N = Number of observations

We analysed this trade-off for the lowest level of coverage i.e., 25%. Observing 25% of animals over 91 days with a fix interval of 16 hours resulted in 136 recorded observations. In contrast, a 365-day sample with 2 hour between fixes yielded 4380 recorded observations. Furthermore, several samples with various settings produced approximately the same amount of observations. For example, the three samples obtained by (0.25, 4, 91), (0.25, 8, 182) and (0.25, 16, 365) configurations generated 546 to 548 observations. Therefore, we used our data to examine what the best strategy is if we have a constraint on the number of observations that may be captured.

Thirdly, we analysed how the accuracy of network metrics is affected when the sampling is done randomly as compared to when it is done group-wise. So for this analysis, we compared the error and the correlation coefficient when the coverage was random versus when whole sets of individuals were sampled. We performed this analysis for 25% and 50% of sampling coverage. As for higher coverage proportions, the two sampling techniques would lead to a similar selection of animals and the differences would be negligible.

## Results

### Error in the values of global network metrics

Error in edge density was most affected by sampling duration and sampling frequency, and least affected by sampling coverage (figure 4). The lowest error was found at the highest sampling duration and highest sampling frequency, even when the coverage was only 25%.

**Figure 4.**
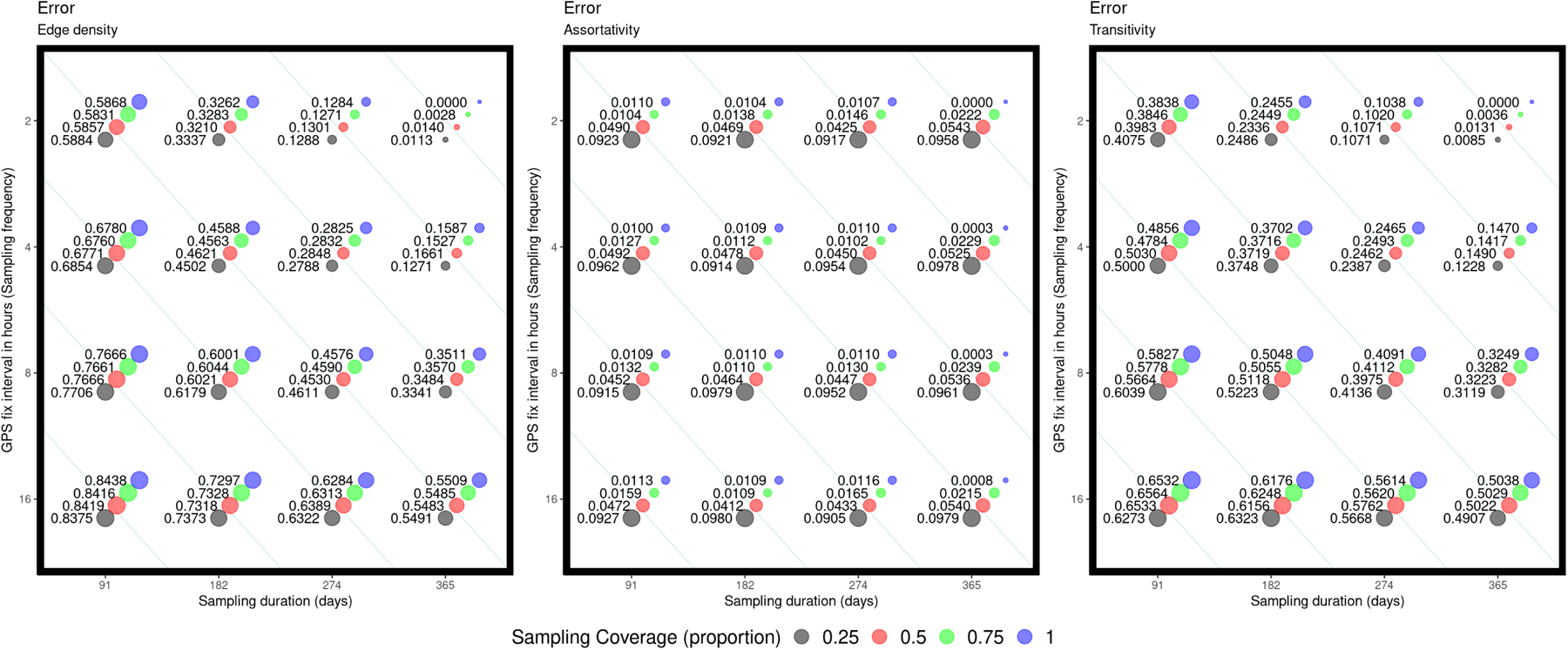
The error in the three global network metrics - edge density, assortativity, and transitivity. The x-axis denotes sampling duration, y-axis denotes fix interval in hours and the colour of the circles in each plot correspond to a sampling coverage value. The size of the circles correspond to the magnitude of the error. Note that the top right value in each plot is zero as it corresponds to the observed network obtained from complete observations of the population.

Overall, assortativity had little error across whole of sampling space (figure 4). We did find error to be slightly elevated with lower sampling coverage (from 0.0113 at 100% coverage and lowest values for duration and frequency to 0.0222 at 75% coverage and highest frequency and duration). However, these errors remain very small, suggesting that assortativity is highly robust to subsampling.

Error in transitivity was significantly influenced by sampling frequency, followed by sampling duration and was least affected by sampling coverage (figure 4). The error rose over 0.5 when the sampling fix interval was increased to 16 hours from 2 hours. However, the error was just 0.008 when the coverage was dropped to 25%. At each level of frequency and duration, the error remained approximately the same regardless of the coverage values. This analysis suggested that the best combination of coverage, frequency and duration depends on the specific network metric being investigated in relation to the research question.

### Correlation coefficient values for node-level network metrics

The network metric degree was most affected by sampling duration, followed by sampling coverage and was least affected by sampling frequency. The rank correlation values dropped below 0.4 when the sampling duration was reduced to 91 days even with full coverage and highest frequency. In contrast, the rank correlation was over 0.7 with a high sampling duration of 365 days and a full coverage, regardless of the observation frequency. Therefore, to get high rank correlation for the network metric degree, higher sampling duration was a critical factor.

Sample coverage had a significant impact on correlation values for network metric strength, followed by sampling duration. After observing individuals for 365 days with a high sampling frequency of 2 hours, the correlation coefficient was as low as 0.39 when just 25% of the population was sampled. In comparison, the correlation coefficient was over 0.75 when the sampling duration was merely 182 days, fix interval was 16 hours, and coverage was 100%. Hence, selecting higher values for coverage and duration was crucial for capturing rank correlations in strength.

The network metric betweenness was least affected by sampling coverage. However, the rank correlation values dropped quickly as the values were lowered on each of the three axes (figure 5). A sampling duration of less than 274 days, fix interval of over 2 hours and coverage less than 50% lead to rank correlations dropping below 0.7. The network metric of betweenness demonstrated the greatest sensitivity to incomplete sampling.

### Trade offs between the three axes of sampling

For the network metrics edge density and transitivity, we found that for a given number of samples, it was typically better to increase sampling duration at the cost of sampling frequency as opposed to increasing sampling frequency at the cost of duration (figure 6a). For example, there were three configurations, resulting in a log (number of observations) of 6.3. Out of these three configurations, for 25% coverage, the lowest error was achieved when the fix interval was 16 hours and a sampling duration of 365 days. The overall error in assortativity remained minimal, less than 0.1 across all configurations. Nevertheless, it was noted that favoring an increase in sampling frequency over sampling duration yielded better results for a fixed number of samples.

While prioritizing sampling duration over sampling frequency provided better rank correlation for the individual-level network metric degree with a fixed number of samples, no distinct trend was evident for the network metrics strength and betweenness (figure 6b). The rank correlation values were under 0.7 for all configurations at 25% sampling coverage for strength and betweenness, indicating the significance of higher sampling coverage in achieving improved rank correlations for these two network metrics.

### Random Vs Group-wise sampling

For the global network metrics, edge density, assortativity and transitivity, the difference in the error as a function of deployment strategy was largely imperceptible (see figure 10 in appendix section Plots for random versus group-wise sampling). However, a close inspection indicated that the error magnitude were mostly lower when the sampling was done groupwise for the network metrics edge density and transitivity. In contrast, random sampling mostly produced a lower error for network measure assortativity.

**Figure 5.**
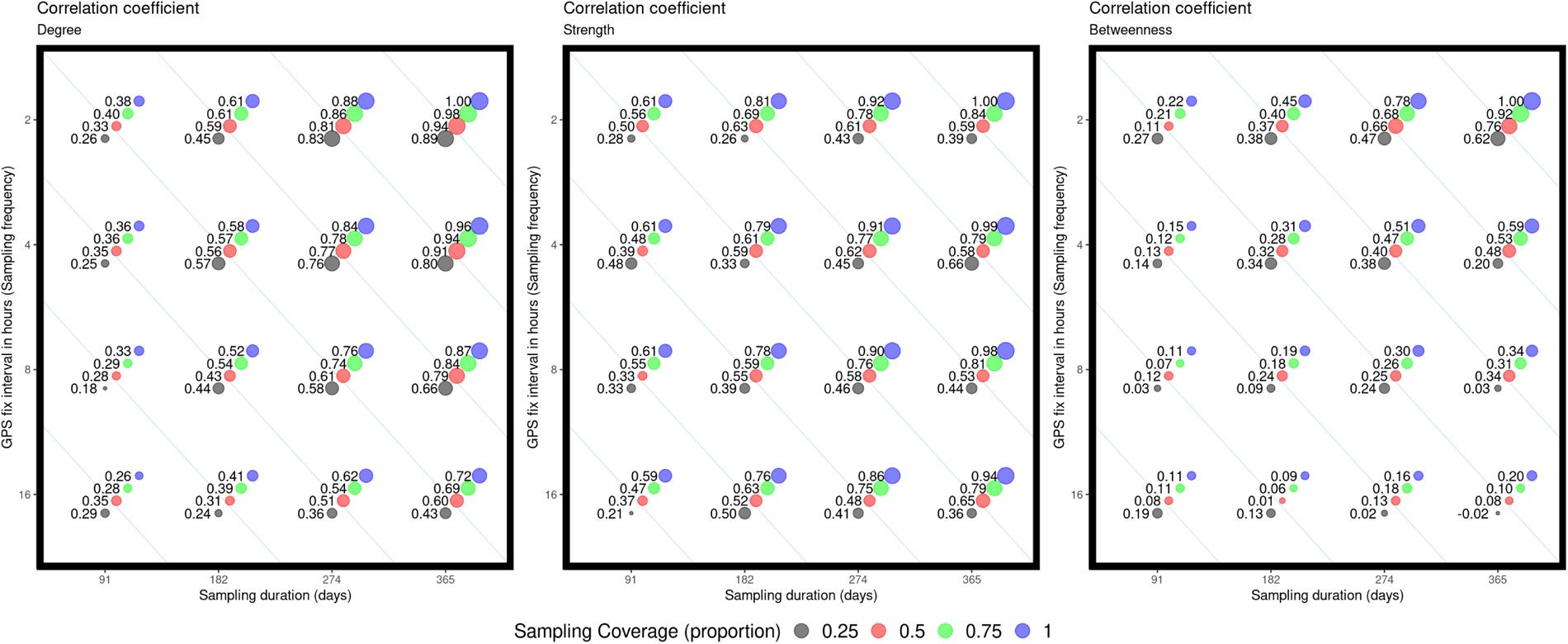
The correlation coefficient between node-level metric values from sampled networks and the fully observed network based on complete observations of the full population. The x-axis represents sampling duration, the y-axis represents fix interval in hours, and the colour of the circles in each plot corresponds to the sampling coverage value. The size of the circles represents the magnitude of the correlation. Note that the top right value in each plot is one as it corresponds to the correlation for the observed network obtained from full observations of the population.

**Figure 6.**
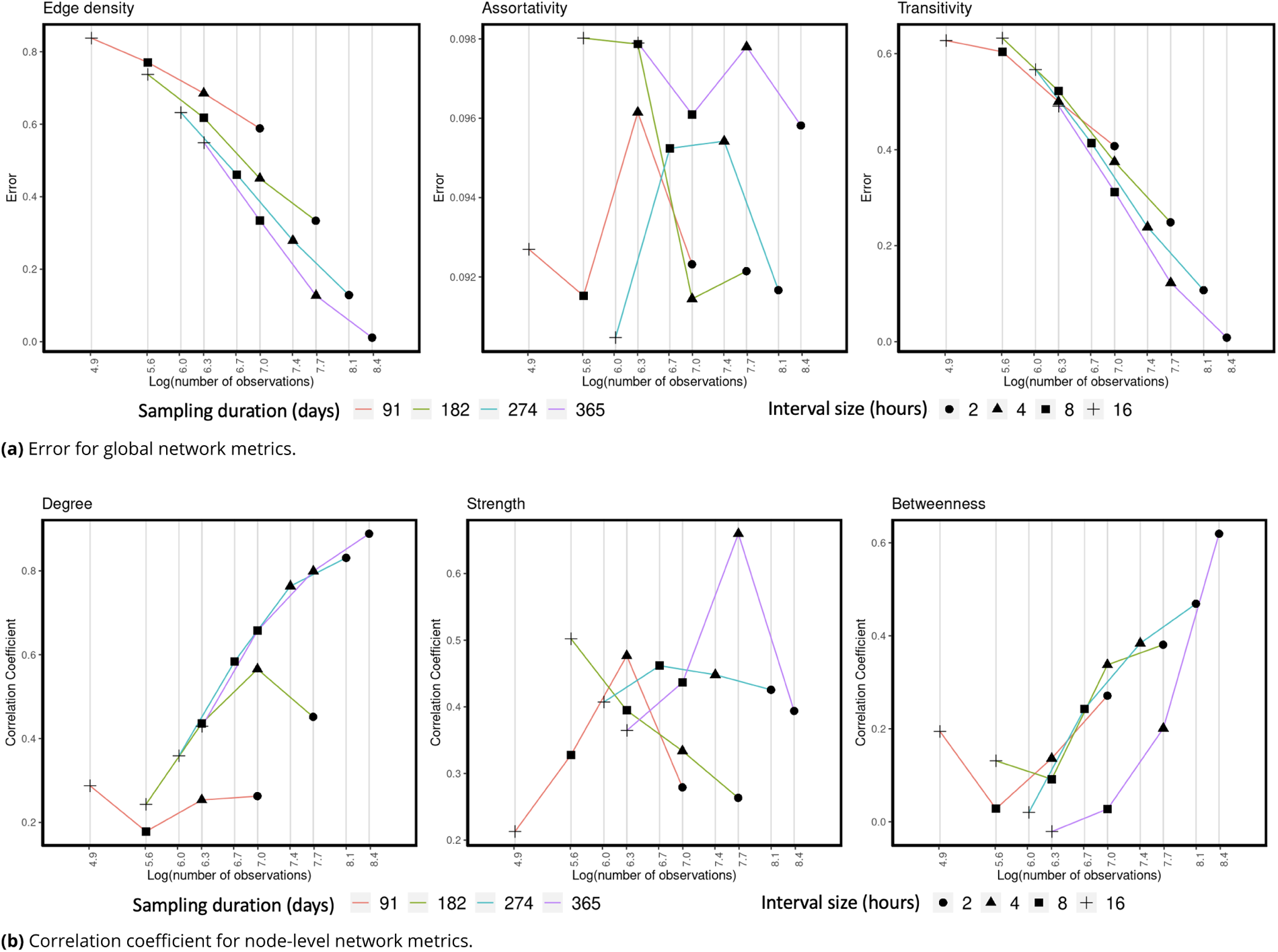
Trade offs in the two axes of sampling, sampling duration and sampling frequency. In each plot, the x-axis denotes the log of total number of observations in the sample configuration. The y-axis denotes the values for error in (a) and correlation coefficients in (b). The color of the lines connecting the points represent sampling duration in days and the shape of the points represent fix interval size in hours.

The differences in the rank correlation values were comparatively more apparent for the individual-level network metrics degree, strength and betweenness (figure 11 in appendix section Plots for random versus group-wise sampling). Degree values tend to correlate better when the sampling is done randomly and the network metric strength correlated better when the sampling was performed group-wise for lower coverage proportions. This held true for nearly all of the 32 samples. Betweenness values for groupwise sampling procedure declined as the sampling efforts were reduced along each of the three sampling axes. This is analogous to the case in which sampling was done at random. However, random sampling yielded better rank correlation for lowest sampling coverage with higher frequency and longer duration.

## Discussion

Our agent-based model, combined with strategic subsampling, highlighted that sampling decisions can have a substantial impact on data collecting results, and hence the suitability of a data set for answering questions about group and individual behaviour (***Davis et al., 2018***; ***Fragaszy et al., 1992***; ***Gilbertson et al., 2021***; ***He et al., 2022***). Our findings suggested that the influence of each of these three axes varies depending on the social network metric, with some metrics requiring more sampling coverage than others to obtain comparable levels of accuracy. We discuss how each of the sampling axes influences the accuracy of estimated network metrics based on the network properties that they represent, and provide recommendations for efficient configurations for the three sampling axes.

### Optimizing sampling strategies for accurate assessment of social network metrics

We examined three aspects of the global network—edge density, assortativity, and transitivity—as well as three aspects of individual-level rankings within the population—degree, strength, and betweenness. Our goal was to identify optimal configurations for the three sampling axes values to minimize error and maximize rank correlations while keeping a fixed number of samples. We found that the global network metrics of edge density and transitivity were significantly impacted by sampling frequency and duration, whereas assortativity was overall much less sensitive (and mostly impacted by sampling coverage). At the node level, both degree and strength were particularly affected by decreasing sampling duration. Sampling coverage had a pronounced effect on strength and betweenness metrics compared to degree. Interestingly, betweenness was heavily influenced by all three sampling axes values.

High sampling coverage is an important factor while calculating such network metrics whose values are not just dependent on a localised part of the network but are influenced by the network individuals as a whole for example, assortativity, and betweenness. Some other network metrics falling into this category are eigenvector centrality, modularity, homophily and closeness centrality. These results are in-line with the conclusions made by ***Davis et al. (2018)*** and Gilbertson et al. (2021) that such measures of association were more sensitive to insufficient sampling. Sampling coverage would also play a crucial role while collecting the samples for research objectives consisting of community detection algorithms in the network. In such research studies, if the sampling coverage is insufficient, observing animals for prolonged periods of time with high frequency will have no positive effect on the estimated values of social network metrics.

High sampling duration and high frequency appears to be necessary for obtaining low error rates in network metrics that are highly dependent on the localised regions of a network, for example, edge density, transitivity, strength, and degree. The accuracy of these network metrics is directly proportional to the comprehensiveness of the monitoring regimes. As a result, monitoring animals with high frequency over an extended period helps to reduce error rates. However, these network metrics are resilient to low sampling coverage since they do not capture information on overall connectivity patterns or clustering within the network, as well as a node’s centrality and influence. These findings align with the study by ***Silk et al. (2015)***, which found that tracking as few as 30% of individuals can still accurately represent their relative positions in the social network, suggesting that collecting more samples per dyad should be prioritised over collecting more individuals (see also ***Davis et al. (2018)***).

### Trade-offs in the three sampling axes

To attain a comparable level of accuracy for edge density and transitivity, lowering the sampling frequency required increasing the sampling duration and coverage while keeping the total number of observations constant. To acquire an error value less than 0.2, more sampling effort was required, as low error values were only attained with a large number of total observations. However, for assortativity, achieving low error rates did not require a large number of observations.

Previous analyses suggested the impact of better sampling duration on the network metrics degree and strength (***Silk et al., 2015***; ***Davis et al., 2018***; ***Voelkl et al., 2011***). Yet ’better sampling’ can be achieved via any of the three axes (duration, frequency, and coverage). Having a high frequency and high coverage at the same time was not required to produce satisfactory values for rank correlation coefficients; rather, at least one of these axes should be high. High frequency sampling was the primary need for achieving favourable values for betweenness, resulting in a large number of observations.

Overall, these findings underscored that it is possible to obtain a desired level of accuracy in the network metric estimates with least amount of sampling effort, if the sampling strategies and the configurations of the three sampling axes are carefully chosen. This choice should always be derived by the properties of the chosen social network metrics along with a careful consideration of which aspects of the network do they capture. However, a challenge is that different axes are better for different network metrics, so which metric should be prioritised will often come down to the research question.

### Optimizing sampling coverage : Insights from random and groupwise coverage

We analysed the differences in the network metric estimates for random and groupwise sampling for lower levels of sampling coverage of 25% and 50%. The network metric assortativity was better estimated for random sampling coverage in contrast to the rest of the two network metrics. This was logical, as assortativity requires both within-type and between-type edges. Therefore, if the research objective centers around assessing assortativity, then the sampling approach should prioritize capturing a representative sample of individuals across various types (e.g. social group, as what we simulated here, or according to traits ***Farine (2014)***).

The node-level network metrics degree and strength showed opposite behaviours to each other with degree performing better with random sampling coverage. A striking difference in the patterns for degree and strength indicates the importance of groupwise connections that contribute significantly towards the total strength of a node in the network. The strength of a node represents the overall intensity and frequency of interactions that a pair of nodes have within the network, where degree just represents the number of individuals a node encounters without considering the intensity of those encounters. Therefore, degree rankings were better preserved in randomly selected network, whereas preserving strength rankings of individuals required more individuals with strong inter-individual preferences. Another way of putting this is that having fewer preferred associates marked will substantially impact an individual’s strength estimate. Therefore, to maximise the accuracy of strength requires sampling sets of more strongly interconnected individuals. Surprisingly, no such pattern was observed for the network metric betweenness, indicating that substantial sampling coverage is necessary for betweenness and that for low levels, it makes no difference whether the coverage is random or groupwise.

This has once again demonstrated the significance of having a well-planned sampling approach that is consistent with the research question and the relevant network metric. The prior selection of a social network metric should guide the sampling coverage criteria.

### Limitations and future directions

Using an agent based model for this analysis has provided us with a lot of flexibility to design a constrained and controlled environment allowing us to compare several scenarios and analyse the effect of sampling levels across the three axes. It has allowed us to capture individual and environmental heterogeneity by modeling agent characteristics, for example, their energy levels and change in state that directly influenced their individual decision-making processes. In addition, it has made it possible to simulate social contacts efficiently using precise principles. This has enabled us to investigate the complex social phenomena and network features that emerge when animals are attracted to, and move with, preferred conspecifics, revealing insights into the fundamental group structures that evolve from bottom-up interactions. Agent-based modelling has also served as a computational platform for hypothesis testing in this paper, allowing us to manipulate model parameters and experiment with different thresholds to test hypotheses about different sampling regimes and the effects of changes in sampling on the accuracy of measured network metrics. This has in turn enabled us to identify critical thresholds that influence the social dynamics and the inferences that we obtain by forming a network structure from the sample.

Following this approach has the potential to greatly inform empirical studies. It is possible to develop testable predictions and theoretical frameworks for studying an animal population (***van der Vaart and Verbrugge, 2008***; ***Franks et al., 2008***; ***Bridge et al., 2017***). The ABM approach has practical uses in conservation and management activities, such as simulating the effects of management interventions and habitat modification on social network structure, stability, and resilience to guide decision-making processes ***Marshall and Duthie (2022)***; ***Koda et al. (2020)***. It can also assist forecast ecological outcomes and population-level effects of social network dynamics. By combining social network data with demographic models, researchers may predict how changes in social structure, connectivity, or network topology will affect population dynamics, dispersal patterns, disease transmission, and conservation outcomes (***Khazaii, 2016***; ***Salgado and Gilbert, 2013***; McLane et al., 2011; ***Ellers et al., 2010***). Further, while simulation models at a conceptual level may not fully capture the complexity of real animal behavior (***Jonker and Treur, 2004***), they can be refined with more specific on-the-ground knowledge from a given population (which itself can be informed from tracking data). Thus, the ABM presented here can be made more realistic by adding additional layers of movement constraints and decision making processes at the level of individuals in an iterative fashion (***Tang and Bennett, 2010***; ***Bernardi and Scianna, 2020***; ***Cristiani et al., 2009***; ***Gochanour et al., 2023***; ***Thiele et al., 2011***).

An important feature of agent-based models is in providing meaningful environmental layers.

Adding environmental barriers, such as impassable terrain or physical hurdles, can be used to test certain hypotheses (***He et al., 2019***). Resource availability can be made more flexible by programming it to include multiple sorts, such as food, and water, whose availability varies across the simulated environment. At the individual level, their behaviours can be adjusted and developed to include more complicated elements that influence their decision-making processes, foraging methods, social dominance, metabolic rates, and other characteristics (***Anderson et al., 2017***). Individuals may also have factors such as stress, reproductive status, and changing preferences for particular types of social affiliations. Another complicated component might be integrating learning processes that allow individuals to adjust their behaviours based on previous experiences, and environmental feedback. For example, the agent based model presented by ***Perez et al. (2021)*** simulated animal movements that were governed by learning and memory in a dynamic landscape and provided insights into how bears use previously acquired landscape data to maximize use of high-quality areas within a heterogeneous landscape. Therefore, by thoroughly considering the features, behaviours, and decision-making processes of species agents in an ABM, researchers may develop realistic simulations that reflect the dynamics of animal behaviour, ecology, and population dynamics within complicated settings (***Hoegh et al., 2021***; de Almeida et al., 2010; ***Diaz et al., 2021***). ***Murphy et al. (2020)*** have presented agent-based modelling as an open-source, accessible, and inclusive tool to serve as a substitute when field data collection is not feasible.

## Conclusions

In conclusion, our research has highlighted the importance of carefully designing sampling strategies aligned with the research question and the specific network metrics relating to the research objectives. The choice of social network metric should guide the criteria for sampling coverage, frequency, and duration, ensuring that the collected data are suitable for addressing questions about the social network structure of the concerned population. Our findings revealed the varying influence of each sampling axis on different network metrics, highlighting the need for tailored sampling approaches to achieve accurate metric estimates.

Moreover, our analysis provided valuable insights into the trade-offs involved in optimizing sampling strategies for social network metric estimation. We identified configurations that yielded minimum error and maximum rank correlations while constraining on the total number of observations. In particular, the sensitivity of network metrics to sampling coverage, frequency, and duration underscores the need for careful consideration of these factors in study design. Our study also highlighted the potential of agent-based modeling as a powerful tool for hypothesis testing and exploration of various scenarios in animal social network analysis. By simulating complex social interactions and integrating environmental constraints, ABM offers a flexible framework for investigating the dynamics of animal behavior and network structure.

Overall, our study contributed to advancing our understanding of how to design studies to collect data with sampling configurations that can maximise the contribution of research effort (time and costs) in terms of producing accurate and reliable animal social networks. Our work underscored the importance of informed sampling strategies and computational modeling approaches in understanding the complexities of animal behavior and network dynamics.

## Acknowledgments

- This publication has emanated from research conducted with the financial support of Science Foundation Ireland under Grant number 18/CRT/6049. For the purpose of Open Access, the author has applied a CC BY public copyright licence to any Author Accepted Manuscript version arising from this submission.
- The work benefited from funding from an Eccellenza Professorship Grant of the Swiss National Science Foundation (Grant Number PCEFP3_187058) and a grant from the European Research Council (ERC) under the European Union’s Horizon 2020 research and innovation programme (grant agreement No. 850859) awarded to Damien Farine.
- P.K. would like to thank the SFI Centre for Research Training in Foundations of Data Science for sponsoring the research visit to the University of Zurich. The assistance was instrumental in facilitating this collaboration.
- P.K. is grateful to the University of Zurich for facilitating her stay and research collaboration during the research visit. The support and resources provided by the Department of Evolutionary Biology and Environmental Studies greatly contributed to the success of this project.

## Appendix

### Model assumptions

1. Each animal is represented as an agent that is capable of making decisions and interacting with it’s environment based on internal states and external stimuli.
2. The grid space is inhabited by a single type of agent.
3. Interactions between animals primarily occur within a local neighborhood defined by the grid structure.
4. Resources required for survival are limited and unevenly distributed across the environment.
5. The resource quantity or quality is not affected by the time of the year. It remains constant throughout the whole cycle of simulations.
6. Agents do not reproduce or exhibit any life history traits.
7. Agent movement patterns are uniform across the simulations, i.e there is no concept of day and night.

### Decision making process of an individual

Figure 7 shows a summary of the decision-making process.

**Appendix 0 Figure 7.**
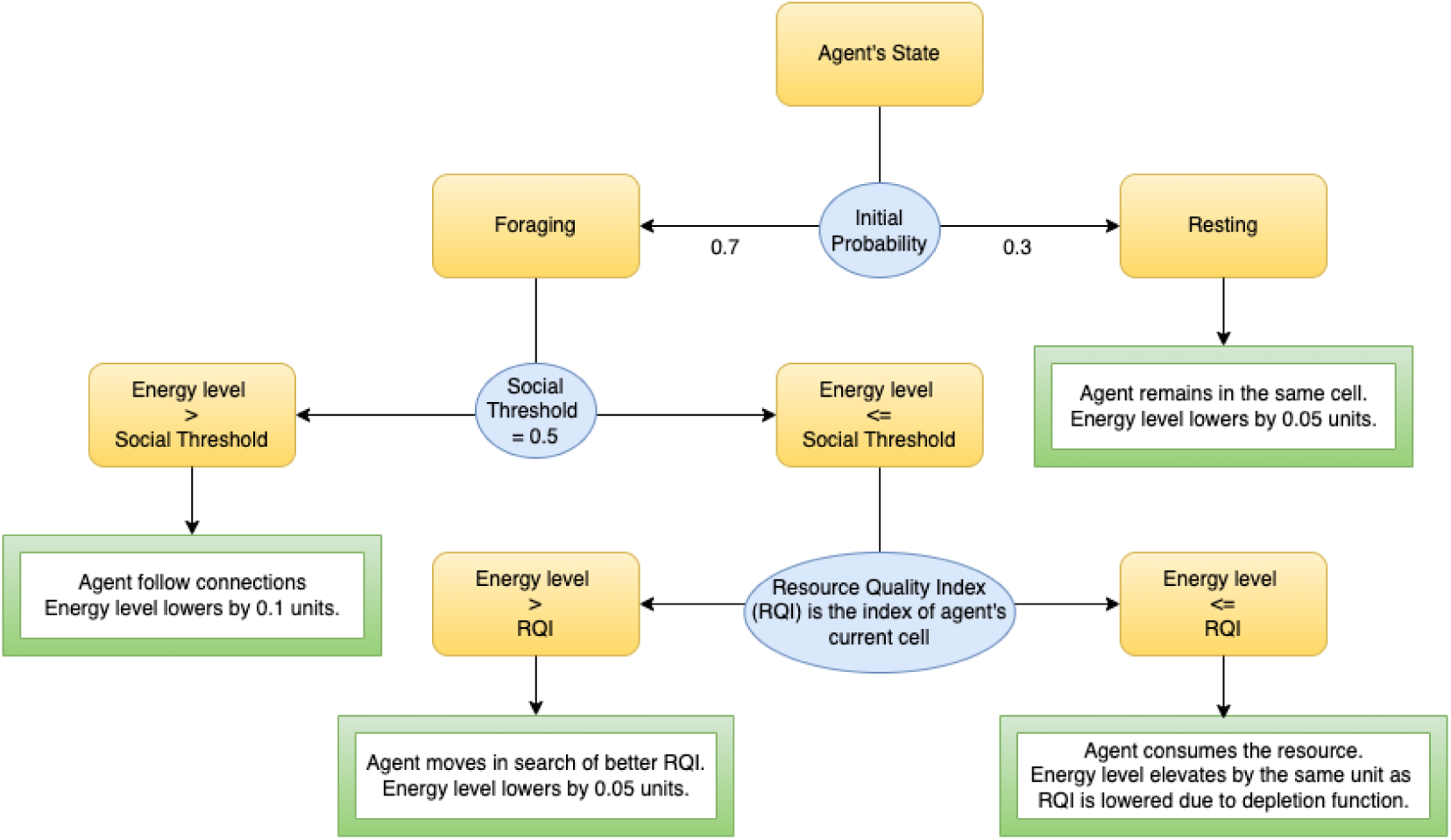
A flowchart depicting an agent’s movement decisions at each time point.

### Agent’s movement when multiple connections are present in multiple neighbouring cells

The agents were designed to undertake a two-step decision process to decide where their next move should be (figure 8).

1. Step 1 : In the first step of this process, agents take an initial decision to move and choose a cell for movement based on the presence of their connections in neighbouring cells. If the connections were present in multiple neighbouring cells, agents decided to move randomly towards one of them. At this step, the agents do not make a move yet but just take an initial decision at this point. All the agents in the simulation decide simultaneously their initial choices but do not move just yet.
2. Step 2 : In step two of this process, it’s time to make a final movement. The final movement decision is based on the choices made by the agents in the first step. For an agent (say A) in the simulation, it’s connections are considered whose initial decision would land them within the immediate neighbourhood of agent A. We then took the mean of all those cells and that rounded up mean value becomes the location of agent A’s final move.

**Appendix 0 Figure 8.**
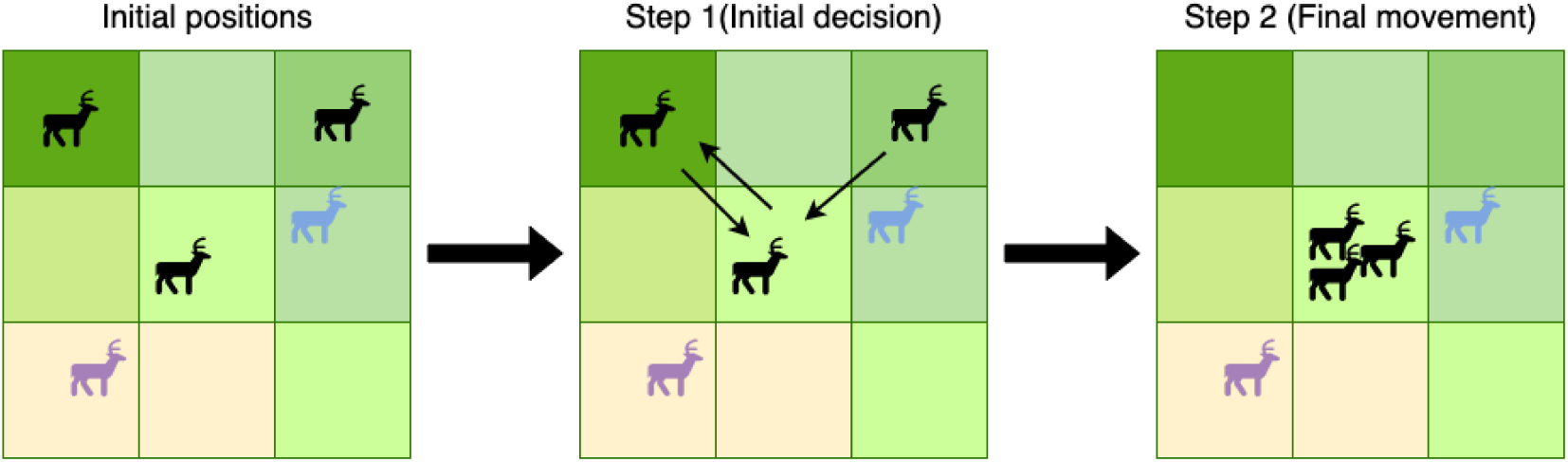
Agents’ two-step decision process when multiple connections are present in the multiple cells of the neighbourhood. The black agents belong to the same group and are all in foraging state with energy levels greater than the social threshold. Their movement decisions are influenced by the presence of their group members in the neighbourhood. In the first step of the process, agents take a decision to move influenced by the presence of their connections but do not make any movement yet. In the second step of the process, they make a collaborative decision. Note that the movement decisions of agents in the black group are not influenced by the presence of other group members in the neighbourhood.

This two-step decision process is performed simultaneously for each agent in the simulation at each time point. The decision process of an agent in foraging state with energy level greater than the social threshold is decided in this way. The decisions are influenced by the presence of all of their connections in the neighbourhood, regardless of their state.

### Model calibration

In the simulations, we calibrated multiple parameters based on agent and environmental properties such that a balance was maintained among the resource availability, agents’ energy levels and their state. For example, the minimum resting threshold was chosen such that it maintained a constant ratio of agents in the foraging and resting state at each timepoint (Figure 9a). Figure 9 depicts the variations in the energy level of an agent at each time point. This value was influenced by the state of the agent as well as the land resource quality index of agent’s current position at each timepoint.

1. Replenish value : Amount by which the resources got replenished at each time point. This value was a function of the current resource value of each cell, total number of agents present in the simulation as well as the size of the grid. The value was defined as :

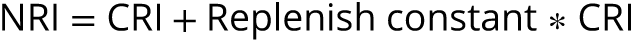

where,

NRI = New resource index,

CRI = Current resource index,

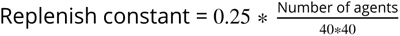

1. Social threshold : It referred to the energy level at which agents prioritize seeking better resources over following their connections. In this context, the social threshold value was set at 0.5.
2. Energy depleted when moving : The quantity of energy expended during movement without resource consumption. This value was 0.1.
3. Energy depleted when resting : Amount of energy depleted while resting. This value was 0.05.
4. Minimum resting threshold : It referred to the minimum energy level at which the agent would choose to forage rather than continue resting. In this instance, the value was set at 0.75.
5. Searching span : It referred to the distance within which agents can search for their connections and subsequently follow them in a given direction. In this context, the search span was configured to extend 2 cells in each direction.
6. Movement span : Number of cells the agents can move at each time point. This value was set to one.

**Appendix 0 Figure 9.**
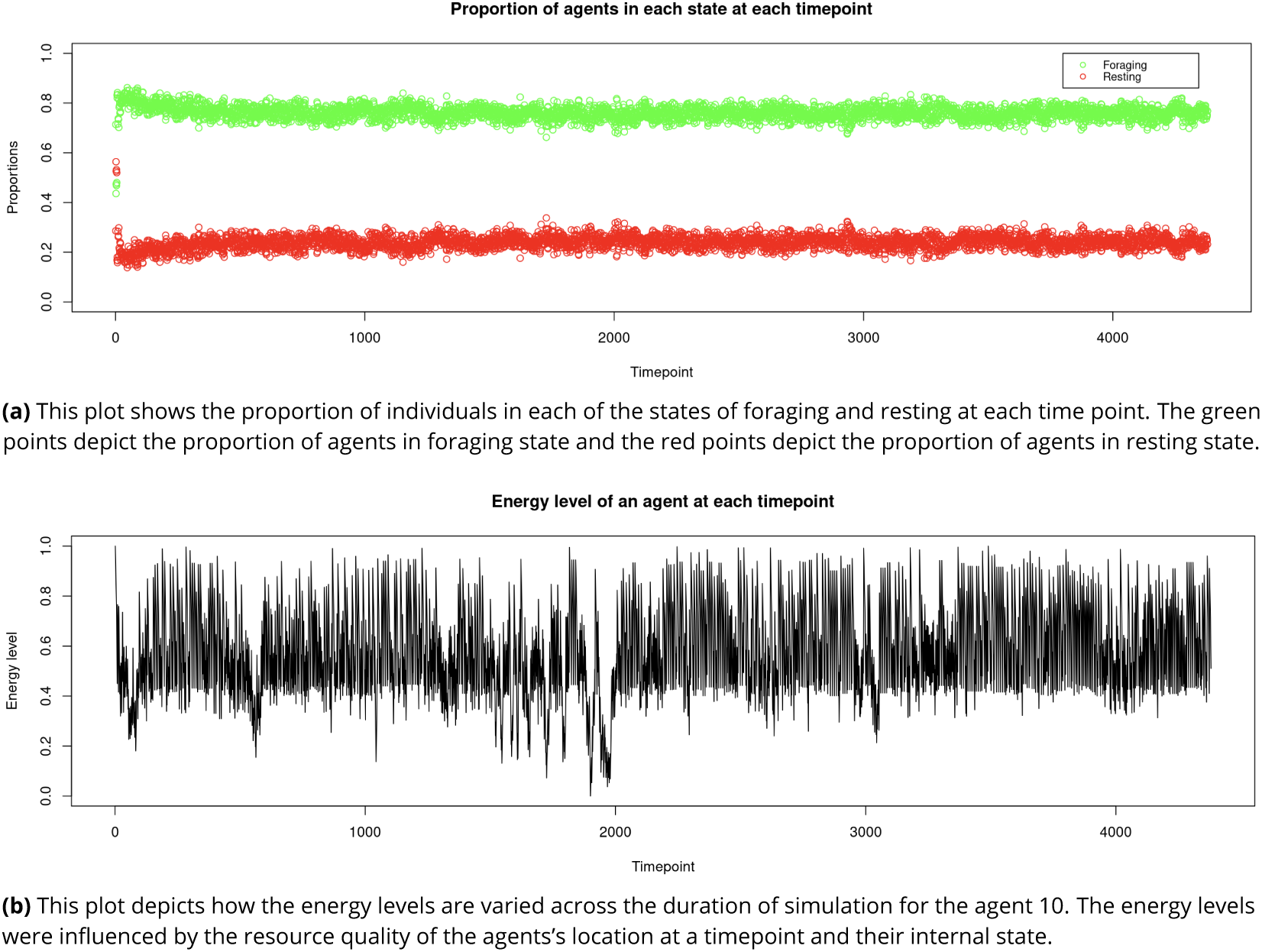
Plots used for model caliberation

## Plots for random versus group-wise sampling

**Appendix 0 Figure 10.**
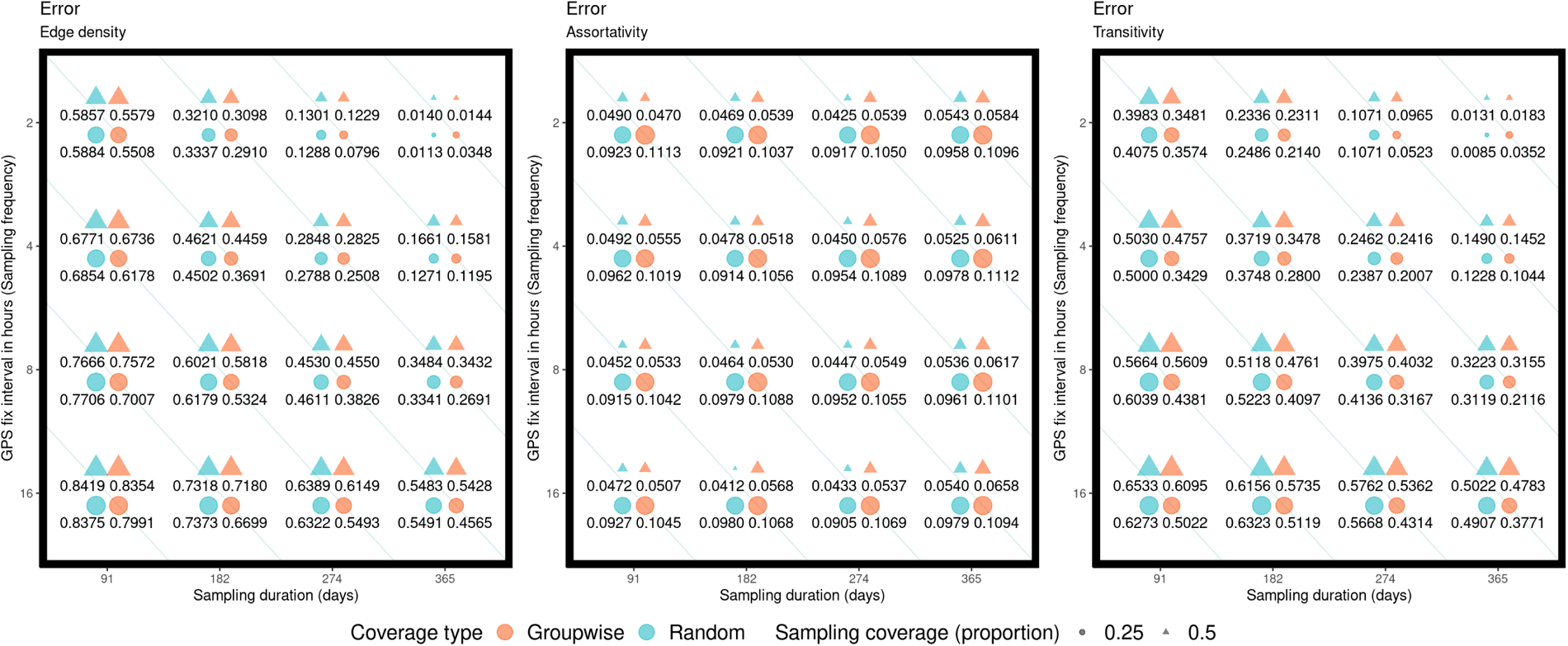
Error comparison between networks resulting from random and groupwise sampling procedures. In each plot, the x-axis represents the sampling duration in days, while the y-axis represents fix rate in hours. Corresponding to each value of the duration and fix rate, the sets of four points reflect the error in network metric values, with the size of each point corresponding to the magnitude of the error. The circular and triangle points indicate the sampling coverage of 25% and 50%, respectively. The black and red coloured points indicate random and groupwise sampling, respectively.

**Appendix 0 Figure 11.**
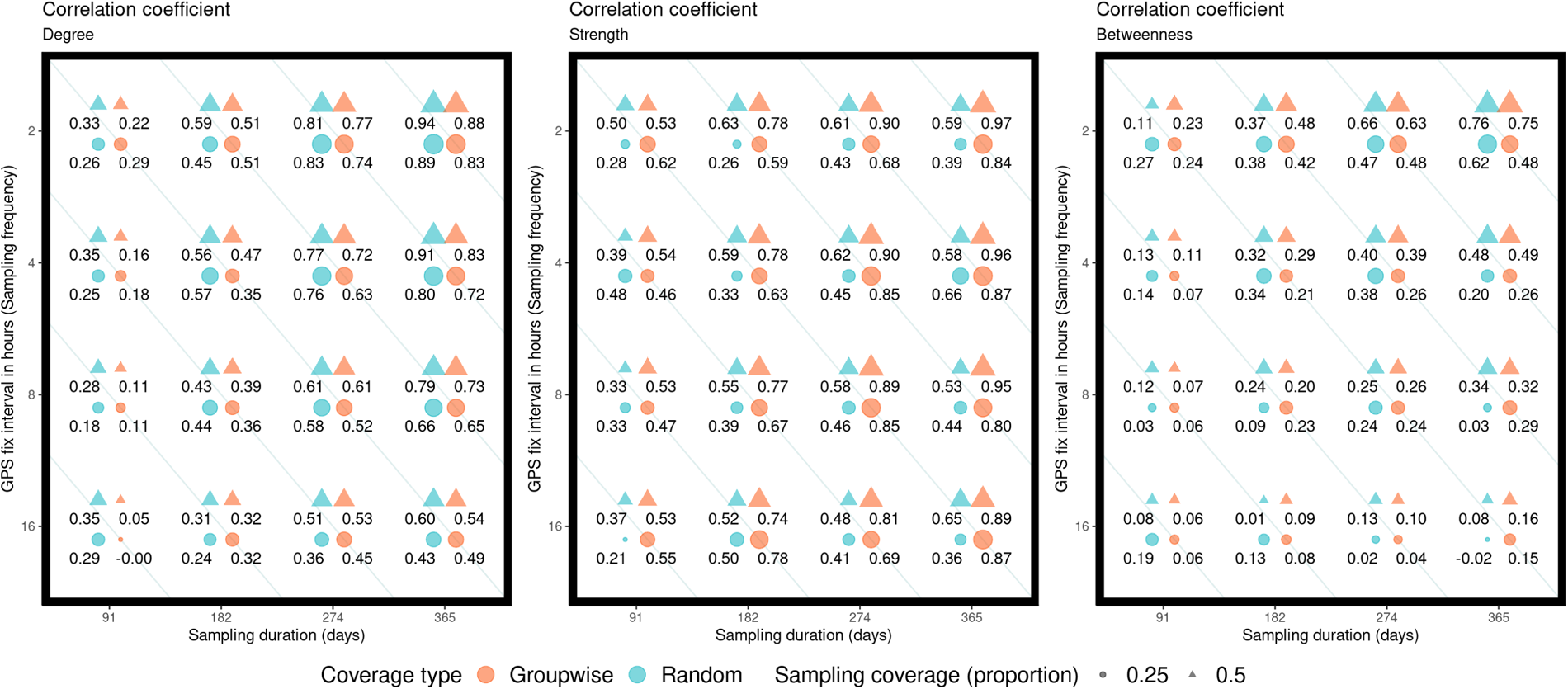
Comparison of correlation coefficient values between networks resulting from random and groupwise sampling procedures. In each plot, the x-axis represents the sampling duration in days and the y-axis represents fix rate in hours. Corresponding to each value of the duration and fix rate, the sets of four points represent the correlation coefficient in network metric values, with the size of each point corresponding to the magnitude of the correlation. The circular and triangle points indicate the sampling coverage of 25% and 50%, respectively. The black and red coloured points indicate random and groupwise sampling, respectively.

## References

Alarcón-Nieto G, Graving JM, Klarevas-Irby JA, Maldonado-Chaparro AA, Mueller I, Farine D. An automated barcode tracking system for behavioural studies in birds. Methods in ecology and evolution. 2018 4; 9(6):1536–1547. https://doi.org/10.1111/2041-210x, doi: 10.1111/2041-210x.13005.

de Almeida SJ, Ferreira RPM, Álvaro E Eiras, Obermayr RP, Geier M, Multi-agent modeling and simulation of an Aedes aegypti mosquito population; 2010. doi: 10.1016/j.envsoft.2010.04.021.

Anderson JH, Downs JA, Loraamm R, Reader S. Agent-based simulation of Muscovy duck movements using observed habitat transition and distance frequencies. Computers, Environment and Urban Systems. 2017 1; 61:49–55. doi: 10.1016/j.compenvurbsys.2016.09.002.

Aplin LM, Firth JA, Farine DR, Voelkl B, Crates R, Čulina A, Garroway CJ, Hinde CA, Kidd L, Psorakis I, Milligan ND, Radersma R, Verhelst B, Sheldon BC. Consistent individual differences in the social phenotypes of wild great tits, Parus major. Animal behaviour. 2015 10; 108:117–127. https://doi.org/10.1016/j.anbehav.2015.07.016, 10.1016/j.anbehav.2015.07.016.

Aureli F, Schaffner CM, Boesch C, Bearder SK, Call J, Chapman CA, Connor RC, Di Fiore A, Dunbar R, Henzi SP, Holekamp KE, Korstjens AH, Layton R, Lee PC, Lehmann J, Manson JH, Ramos-Fernández G, Strier KB, Van Schaik CP. Fission-Fusion Dynamics. Current anthropology. 2008 8; 49 (4):627–654. https://doi.org/10.1086/586708, doi: 10.1086/586708.

Bernardi S, Scianna M. An agent-based approach for modelling collective dynamics in animal groups distinguishing individual speed and orientation. Philosophical transactions - Royal Society Biological sciences. 2020 7; 375(1807):20190383. https://doi.org/10.1098/rstb.2019.0383, doi: 10.1098/rstb.2019.0383.

Bridge ES, Ross JD, Contina A, Kelly JF. Using Agent-Based Models to Scale from Individuals to Populations; 2017. https://doi.org/10.1007/978-3-319-68576-2_11 10.1007/978-3-319-68576-2_11.

Costenbader E, Valente TW. The stability of centrality measures when networks are sampled. Social Networks. 2003 10; 25:283–307. doi: 10.1016/S0378-8733(03)00012-1.

Cristiani E, Frasca P, Piccoli B. Effects of anisotropic interactions on the structure of animal groups. . 2009 3; http://arxiv.org/abs/0903.4056.

Croft DP, James R, Krause J. Exploring Animal Social Networks. Princeton University Press; 2008.

Cross PC, Creech TG, Ebinger MR, Heisey DM, Irvine KM, Creel S. Wildlife contact analysis: Emerging methods, questions, and challenges. Behavioral Ecology and Sociobiology. 2012 10; 66:1437–1447. doi: 10.1007/s00265-012-1376-6.

Davis GH, Crofoot MC, Farine DR. Estimating the robustness and uncertainty of animal social networks using different observational methods. Animal Behaviour. 2018 7; 141:29–44. doi: 10.1016/j.anbehav.2018.04.012.

Diaz SG, DeAngelis DL, Gaines MS, Purdon A, Mole MA, van Aarde RJ. Development and validation of a spatially-explicit agent-based model for space utilization by African savanna elephants (Loxodonta africana) based on determinants of movement. Ecological Modelling. 2021 5; 447. doi: 10.1016/j.ecolmodel.2021.109499.

Ellers J, Hoogendoorn M, Wendt D. An agent-based modeling approach to investigate emergent patterns in ecological systems. In:, vol. 2; 2010. p. 6–13. doi: 10.1109/WI-IAT.2010.200.

Espín-Noboa L, Wagner C, Karimi F, Lerman K. Towards Quantifying Sampling Bias in Network Inference. Companion Proceedings of the The Web Conference. 2018 1; https://doi.org/10.1145/3184558.3191567 10.1145/3184558.3191567.

Farine D. assortnet: Calculate the Assortativity Coefficient of Weighted and Binary Networks; 2016, https://CRAN.R-project.org/package=assortnet, rpackage version 0.20.

Farine DR. Measuring phenotypic assortment in animal social networks: Weighted associations are more robust than binary edges. Animal Behaviour. 2014 3; 89:141–153. doi: 10.1016/j.anbehav.2014.01.001.

Farine DR, Whitehead H. Constructing, conducting and interpreting animal social network analysis. Journal of Animal Ecology. 2015 9; 84:1144–1163. doi: 10.1111/1365-2656.12418.

Fragaszy DM, Boinski S, Whipple J. Behavioral sampling in the field: Comparison of individual and group sampling methods. American journal of primatology. 1992 1; 26(4):259–275. https://doi.org/10.1002/ajp.1350260404 doi: 10.1002/ajp.1350260404.

Franks DW, James R, Noble J, Ruxton GD. Developing a methodology for social network sampling. In: IEEE Symposium on Artificial Life; 2008. https://api.semanticscholar.org/CorpusID:37065379.

Fründ J, McCann KS, Williams NM. Sampling bias is a challenge for quantifying specialization and network structure: lessons from a quantitative niche model. Oikos. 2015 7; 125(4):502–513. https://doi.org/10.1111/oik.02256, doi: 10.1111/oik.02256.

Gilbertson MLJ, White LA, Craft ME. Trade-offs with telemetry-derived contact networks for infectious disease studies in wildlife. Methods in Ecology and Evolution. 2021 1; 12:76–87. doi: 10.1111/2041-210X.13355.

Gochanour B, Fernández-López J, Contina A. abmR: An R package for agent-based model analysis of large-scale movements across taxa. Methods in Ecology and Evolution. 2023 1; 14:218–230. doi: 10.1111/2041-210X.14014.

Grimm V, Railsback SF, Vincenot CE, Berger U, Gallagher C, Jiang J, Edmonds B, Ge J, Giske J, Groeneveld J, Johnston ASA, Milles A, Nabe–Nielsen J, Polhill G, Radchuk V, Rohwäder M, Stillman RA, Thiele JC, Ayllón D. The ODD Protocol for Describing Agent-Based and Other Simulation Models: a second update to improve clarity, replication, and structural realism. JASSS. 2020 1; 23(2). https://doi.org/10.18564/jasss.4259, doi: 10.18564/jasss.4259.

He P, Klarevas-Irby JA, Papageorgiou D, Christensen C, Strauss ED, Farine DR. A guide to sampling design for GPS -based studies of animal societies. Methods in Ecology and Evolution. 2022 10; https://onlinelibrary.wiley.10.1111/2041-210X.13999, doi: 10.1111/2041-210X.13999.

He P, Maldonado-Chaparro AA, Farine DR. The role of habitat configuration in shaping social structure: a gap in studies of animal social complexity. Behavioral Ecology and Sociobiology. 2019 1; 73. doi: 10.1007/s00265-018-2602-7.

Hebblewhite M, Haydon DT. Distinguishing technology from biology: a critical review of the use of GPS telemetry data in ecology. Philosophical transactions - Royal Society Biological sciences. 2010 7; 365(1550):2303–2312. 10.1098/rstb.2010.0087, doi: 10.1098/rstb.2010.0087.

Hoegh A, van Manen FT, Haroldson M. Agent-Based Models for Collective Animal Movement: Proximity-Induced State Switching. Journal of Agricultural, Biological, and Environmental Statistics. 2021 12; 26:560–579. doi: 10.1007/s13253-021-00456-0.

Hoppitt WJE, Farine DR. Association indices for quantifying social relationships: how to deal with missing observations of individuals or groups. Animal Behaviour. 2018 2; 136:227–238. doi: 10.1016/j.anbehav.2017.08.029.

James R, Croft DP, Krause J. Potential banana skins in animal social network analysis. . 2009; http://opus.bath.ac.uk/14182/.

Jonker CM, Treur J. Agent-Based Simulation of Animal Behaviour. Applied intelligence. 2004 1; 15(2):83–115. https://doi.org/10.1023/a:1011246304650, doi: 10.1023/a:1011246304650.

Jung TS, Hegel T, Bentzen TW, Egli K, Jessup L, Kienzler M, Kuba K, Kukka PM, Russell K, Suitor MJ, Takagi K. Accuracy and performance of low-feature GPS collars deployed on bison Bison bison and caribou Rangifer tarandus. Wildlife biology. 2018 1; 2018(1):1–11. https://doi.org/10.2981/wlb.00404, doi: 10.2981/wlb.00404.

Kaur P. aniSNA: Statistical Network Analysis of Animal Social Networks; 2024, r package version 1.1.1.

Kaur P, Ciuti S, Ossi F, Cagnacci F, Loison A, Atmeh K, McLoughlin P, Reinking AK, Beck JL, Ortega AC, Kauffman M, Boyce MS, Salter-Townshend M. Assessing bias and robustness of social network metrics using GPS based radio-telemetry data. . 2023; 18. https://doi.org/10.1101/2023.03.30.534779 10.1101/2023.03.30.534779.

Khazaii J. Agent-Based modeling; 2016. https://doi.org/10.1007/978-3-319-33328-1_13, doi: 10.1007/978-3-319-33328-1_13.

Koda H, Arai Z, Matsuda I. Agent-based simulation for reconstructing social structure by observing collective movements with special reference to single-file movement. PloS one. 2020 12; 15(12):e0243173. https://doi.org/10.1371/journal.pone.0243173, doi: 10.1371/journal.pone.0243173.

Krause J, Croft DP, James R. Social network theory in the behavioural sciences: Potential applications. Behavioral Ecology and Sociobiology. 2007; 62(1):15–27. doi: 10.1007/s00265-007-0445-8.

Krause J, Krause S, Arlinghaus R, Psorakis I, Roberts S, Rutz C. Reality mining of animal social systems. Trends in Ecology and Evolution. 2013 9; 28:541–551. doi: 10.1016/j.tree.2013.06.002.

Lusseau D, Whitehead H, Gero S. Incorporating uncertainty into the study of animal social networks. Animal Behaviour. 2008; doi: 10.1016/j.anbehav.2007.10.029.

Marshall BM, Duthie AB. abmAnimalMovement: An R package for simulating animal movement using an agent-based model. F1000Research. 2022 10; 11:1182. doi: 10.12688/f1000research.124810.1.

McLane AJ, Semeniuk C, McDermid GJ, Marceau DJ, The role of agent-based models in wildlife ecology and management; 2011. doi: 10.1016/j.ecolmodel.2011.01.020.

Murphy KJ, Ciuti S, Kane A. An introduction to agent-based models as an accessible surrogate to field-based research and teaching. Ecology and evolution. 2020 10; 10(22):12482–12498. https://doi.org/10.1016/j.anbehav.2010.06.020, doi: 10.1002/ece3.6848.

Perez AZ, Bone C, Stenhouse G. Simulating multi-scale movement decision-making and learning in a large carnivore using agent-based modelling. Ecological Modelling. 2021 7; 452. doi: 10.1016/j.ecolmodel.2021.109568.

Perreault **C**. A note on reconstructing animal social networks from independent small-group observations. Animal behaviour. 2010 9; 80(3):551–562. https://doi.org/10.1016/j.anbehav.2015.07.016, doi: 10.1016/j.anbehav.2010.06.020.

R Core Team. R: A Language and Environment for Statistical Computing. R Foundation for Statistical Computing, Vienna, Austria; 2022, https://www.R-project.org/.

Salgado M, Gilbert N. Agent-Based modelling; 2013. https://doi.org/10.1007/978-94-007-7052-2_4 doi: 10.1007/978-94-007-7052-2_4.

Shizuka D, Barve S, Johnson AE, Walters EL. Constructing social networks from automated telemetry data: A worked example using within- and across-group associations in cooperatively breeding birds. Methods in ecology and evolution. 2021 11; 13(1):133–143. https://doi.org/10.1111/2041-210x.13737, doi: 10.1111/2041-210x.13737.

Silk MJ. The next steps in the study of missing individuals in networks: a comment on Smith et al. (2017). Social Networks. 2018 1; 52:37–41. doi: 10.1016/j.socnet.2017.05.002.

Silk MJ, Jackson AL, Croft DP, Colhoun K, Bearhop S. The consequences of unidentifiable individuals for the analysis of an animal social network. Animal Behaviour. 2015 6; 104:1–11. doi: 10.1016/j.anbehav.2015.03.005.

Smith JA, Moody J. Structural effects of network sampling coverage I: Nodes missing at random. Social Networks. 2013; 35(4):652–668. doi: 10.1016/j.socnet.2013.09.003.

Smith JA, Morgan J. Network Sampling Coverage II: The Effect of Non-random Missing Data on Network Measurement 1 Network Sampling Coverage II: The Effect of Non-random Missing Data on Network Measurement. . 2016; http://www.elsevier.com/open-access/userlicense/1.0/1.

Smith JA, Morgan JH, Moody J. Network sampling coverage III: Imputation of missing network data under different network and missing data conditions. Social Networks. 2022; 68:148–178. doi: 10.1016/j.socnet.2021.05.002.

Sosa S, Jacoby DMP, Lihoreau M, Sueur C. Animal social networks: Towards an integrative framework em-bedding social interactions, space and time. Methods in Ecology and Evolution. 2021 1; 12:4–9. doi: 10.1111/2041-210X.13539.

Strandburg-Peshkin A, Farine D, Couzin ID, Crofoot MC. Shared decision-making drives collective movement in wild baboons. Science. 2015 6; 348(6241):1358–1361. https://doi.org/10.1126/science.aaa5099, doi: 10.1126/science.aaa5099.

Tan RK, Bora S. Adaptive parameter tuning for agent-based modeling and simulation. Simulation. 2019 6; 95(9):771–796. https://doi.org/10.1177/0037549719846366, doi: 10.1177/0037549719846366.

Tang W, Bennett DA. Agent-based Modeling of Animal Movement: A review. Geography compass. 2010 7; 4(7):682–700. https://doi.org/10.1111/j.1749-8198.2010.00337.x, doi: 10.1111/j.1749-8198.2010.00337.x.

Thiele JC, Kurth W, Grimm V. Agent-and Individual-based Modeling with NetLogo: Introduction and new NetLogo Extensions. 2011.

van der Vaart E, Verbrugge R. Agent-based models for animal cognition: a proposal and prototype. In: Adaptive Agents and Multi-Agent Systems; 2008. https://api.semanticscholar.org/CorpusID:6592910.

Voelkl B, Kasper C, Schwäb C. Network measures for dyadic interactions: stability and reliability. American journal of primatology. 2011 3; 73(8):731–740. https://doi.org/10.1002/ajp.20945, doi: 10.1002/ajp.20945.

Wey T, Blumstein DT, Shen W, Jordán F. Social network analysis of animal behaviour: a promising tool for the study of sociality. Animal Behaviour. 2008 2; 75:333–344. doi: 10.1016/j.anbehav.2007.06.020.

